# Early precision of radial patterning of the mouse cochlea is achieved by a linear BMP signaling gradient and is further refined by SOX2

**DOI:** 10.1101/2022.08.30.505910

**Authors:** Matthew J. Thompson, Vidhya Munnamalai, David M. Umulis

## Abstract

Positional information encoded in signaling molecules is essential for early patterning in the prosensory domain of the developing cochlea. The cochlea contains an exquisite repeating pattern of sensory hair cells and supporting cells. This requires precision in the morphogen signals that set the initial radial compartment boundaries, but this has not been investigated. To measure gradient formation and morphogenetic precision in developing cochlea, we developed a quantitative image analysis procedure measuring SOX2 and pSMAD1/5/9 profiles in mouse embryos at embryonic day (E)12.5, E13.5, and E14.5. Intriguingly, we found that the pSMAD1/5/9 profile forms a linear gradient in the medial ∼75% of the PSD during E12.5 and E13.5. This is a surprising activity readout for a diffusive BMP4 ligand secreted from a tightly constrained lateral region^1,2^ since morphogens typically form exponential or power-law gradient shapes. This is meaningful for gradient interpretation because while linear profiles offer the theoretically highest information content and distributed precision for patterning, a linear morphogen gradient has not yet been observed. In addition to the information-optimized linear profile, we found that while pSMAD1/5/9 is stable during this timeframe, an accompanying gradient of SOX2 shifts dynamically. Third, we see through joint decoding maps of pSMAD1/5/9 and SOX2 that there is a high-fidelity mapping between signaling activity and position in the regions soon to become Kölliker’s organ and the organ of Corti, where radial patterns are more intricate than lateral regions. Mapping is ambiguous in the prosensory domain precursory to the outer sulcus, where cell fates are uniform. Altogether, this research provides new insights into the precision of early morphogenetic patterning cues in the radial cochlea prosensory domain.

**Summary Paragraph:** The organ of Corti is the precisely patterned group of cells in the cochlea responsible for transforming sound energy into our perception of hearing. Morphogenetic signals encoding positional information are crucial for the early stages of patterning along the developing cochlea’s radial axis. SOX2 and pSMAD1/5/9 are transcription factors that together serve as an integrative readout of morphogen activity during E12.5 to E14.5 in the developing mouse cochlea. However, the role of spatiotemporal precision in these signals is unknown. Here we show that pSMAD1/5/9 forms a linear profile to establish a domain spanning reference frame of positional information and that SOX2 further refines precision. We found that the pSMAD1/5/9 signal retains its linear shape across at least 24 h of development while SOX2 dynamically shifts. The stable linear pSMAD1/5/9 profile provides a global reference point of radial positional information, while the SOX2 profile improves local precision with steep slopes. Furthermore, a linear profile from a diffusive ligand is unexpected, suggesting unidentified mechanisms of BMP regulation unique to this system. A version of the source-sink model for creating a linear morphogen profile modified from its original formulation^3^ is explored in this system, enabling a tight fit between the BMP model and pSMAD1/5/9 data. We expect the methods and results shown here to be a starting point for increased precision in cochlear morphogen activity measurements to enable further modeling and experimental inquiry. This combination of quantitative mechanistic explanation for how signals form, along with quantitative interpretations of their decoding properties, revealing why they form a certain way, together form a potent basis for biological discovery and may even be applied to the design of synthetic systems.

## Introduction

The mammalian organ of Corti (OC) is a highly specialized sensory epithelium housed in the cochlea of the inner ear. It consists of cell types that are spatially and structurally organized to work together and transform acoustic energy into the electrochemical signals sent up to central auditory pathways to enable hearing. During development, patterning cues coordinate across the radial axis of the OC to subdivide the sensory hair cells (HCs) into two classes: medial sound-detecting inner hair cells (IHCs) along with their associated supporting cells (SCs) and lateral electromotive sound-amplifying outer hair cells (OHCs) along with their associated SCs. This radial pattern repeats along the longitudinal axis that controls frequency selectivity. To achieve this precision in organization and sensory function, the developmental programs guiding the process of dynamic cell fate specification must also be precise. This precision begins with molecular signaling across the radial axis during the early stages of OC development when morphogens establish the positional information crucial for initiating this pattern-formation process^4^.

The OC on the floor of the cochlear duct originates from the prosensory domain (PSD) marked by SOX2 expression at embryonic day (E) 12.5^5^ in the mouse cochlea. This domain develops into the highly ordered sensory organ characterized as follows: In the mature cochlea, the OC is flanked by a non-sensory medial inner sulcus and lateral outer sulcus. In the OC, the HCs are intercalated by SCs, which provide trophic and structural support to the HCs. The mature status of OC and the inner and outer sulci are in place by around postnatal day (P) 12. Altogether, the radial OC pattern consists of 11 HC and SC types. Transcriptomic identities subdivide the seven SC types into medial and lateral sensory subtypes^6^, where medial SCs include inner border cells (IBCs) and inner phalangeal cells (IPhCs), and lateral SCs include inner pillar cells (IPCs), outer pillar cells (OPCs), and three rows of Deiters’ cells. The single row of IHCs is positioned in the medial portion of the OC, and three rows of OHCs are situated in the lateral portion, separated by the tunnel of Corti that is flanked by the pillar cells.

Cochlear development progresses through four successive fate-restricting phases based on the active radial patterning program (**Figure 1**). These phases are nominal, and some overlap exists at their borders. Phase one of cochlear outgrowth occurs from ∼E10.5 to E13 (**Figure 1A**). Here, a prosensory domain marked by widespread SOX2 signaling is established in the floor of the cochlear duct^5^. A JAG1-Notch-mediated lateral induction program sets this domain. Phase two is characterized by a transition from lateral induction to lateral inhibition, which drives the onset of HC differentiation via Delta (DL)-Notch-mediated signaling (**Figure 1B**). This initiates the mosaic organization of HCs and SCs starting around late E14.5 after HC fate specification is initiated by expression of the ATOH1, a HC-specific transcription factor^7^. The precision of signals in phase one establishes the domain landmarks in phase two, where errors could propagate. JAG1 and SOX2 expression in the PSD becomes refined and is restricted to the sensory epithelium. Also, within phase two, beginning around E14.5^8^, a transient group of medial columnar cells emerges comprising Kölliker’s organ (KO). It is a temporary structure essential for the cultivation of neural connections^9^, is a major signaling center during development^10–12^, and contains radially patterned cell types^13^. Phase three of OC patterning occurs through late gestation from ∼E14 to P0. It is characterized by the primary role of mechanically driven refinement to the morphogen-mediated patterns initiated in the previous phases^14–16^ (**Figure 1C**). Lastly, phase four is characterized by the maturation of cell types within their established radial pattern (**Figure 1D**) and apoptotic transformation of KO to form the inner sulcus.

**Figure 1.**
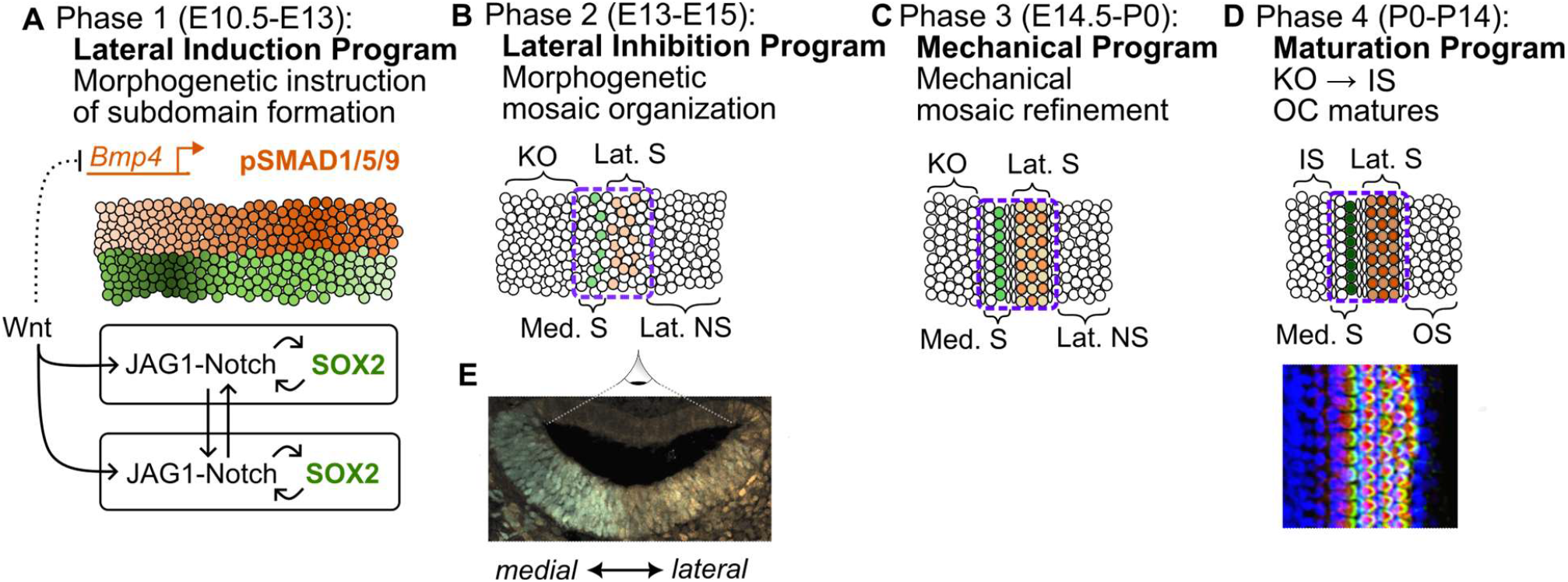
Patterning in the cochlear floor begins with morphogenetic information transmission and is refined mechanically. **(A-D)** top-down views of the floor of the cochlear duct across chronological phases, as depicted in an E12.5 cross-section in **E. (A)** Phase one is characterized by the lateral induction program where position-encoding transcription factors instruct the organization of subdomains that emerge within phase two. The activity of SOX2 and pSMAD1/5/9 is illustrated as the set of two principal readouts of positional information during this phase. For (B-D), green and orange represent medial and lateral HCs rather than SOX2 and pSMAD1/5/9 explicitly. **(B)** During phase two, Delta-Notch lateral inhibition creates a rough mosaic of nascent inner (light green) and outer (light orange) hair cells (HCs) with their associated supporting cells within the primordial organ of Corti (OC; dashed outline). Kölliker’s organ (KO) is established in the medial non-sensory region, which also has radial cell specification (not depicted). **(C)** In late phase two and through phase three, shear stresses and differential adhesion properties between cell types cause a refinement of the OC pattern. **(D)** Phase four, beginning near birth and extending through the onset of hearing, includes the maturation of these domains. KO transitions to the inner sulcus (IS), hair cells and supporting cells in the OC develop and mature within their pattern, and the outer sulcus (OS) forms from the lateral non-sensory region. (D) bottom: a confocal image of a mature cochlea; blue: SOX2, red: Myo6 (HC marker), green: Phalloidin to mark F-actin (stereocilia “hair” marker).

Since cell fate specification patterns begin in phase two, the precision of the morphogenetic signals in phase one determines the precision in these landmark patterns. Positional information readily accessible for measurement and quantification is encoded in transcription factor concentration profiles^17,18^. However, it must be kept in mind that this information is not entirely accessible to cells for ‘decision-making’ since other noisy and context-dependent processes (e.g., promoter accessibility) sit between a transcription factor and its target gene expression^19^. Taking such limitations into account, transcription factor concentration profiles can be seen as an upper limit of the amount of information available for interpretation by cells; an assumption referred to as optimal decoding^18^.

Recent studies on Drosophila segmentation^20^ and the mouse neural tube^21,22^ show that profiles encoding positional information are more reproducible than initially anticipated, influencing the interpretation of developmental programs. While mechanical forces drive the refinement of the precise relative positioning of differentiated cells in the sensory epithelium^14^, the precision of morphogenetic positional information needed to establish earlier subdomains has not yet been investigated.

## Results

### Quantification of Radial Transcription Factor Profiles in the Developing Cochlea

To quantify how positional information is encoded across the radial axis of the cochlear floor, we select SOX2 and phosphorylated SMAD1/5/9 (pSMAD1/5/9) for co-expression analysis at E12.5, E13.5, and E14.5. These signals are active in a radial counter-gradient, where SOX2 is high medially and low laterally, and pSMAD1/5/9 is high laterally and low medially (**Figure 2A-D**) at the timepoints measured. pSMAD1/5/9 represents the active form of the receptor-regulated SMADs (R-SMADs) acting as secondary signal transducers in the bone morphogenetic protein (BMP) pathway. Its contribution is indispensable for radial patterning^1,23,24^. The SOX2 transcription factor is a necessary prosensory signal in the system at this stage^25,26^ and is expressed as an integrating readout of JAG1^27^ and FGF20^28,29^ activity, where *Jag1* and *Fgf20* are predicted Wnt target genes^23,30^. Both transcription factors provide information for cochlear patterning and organization.

**Figure 2.**
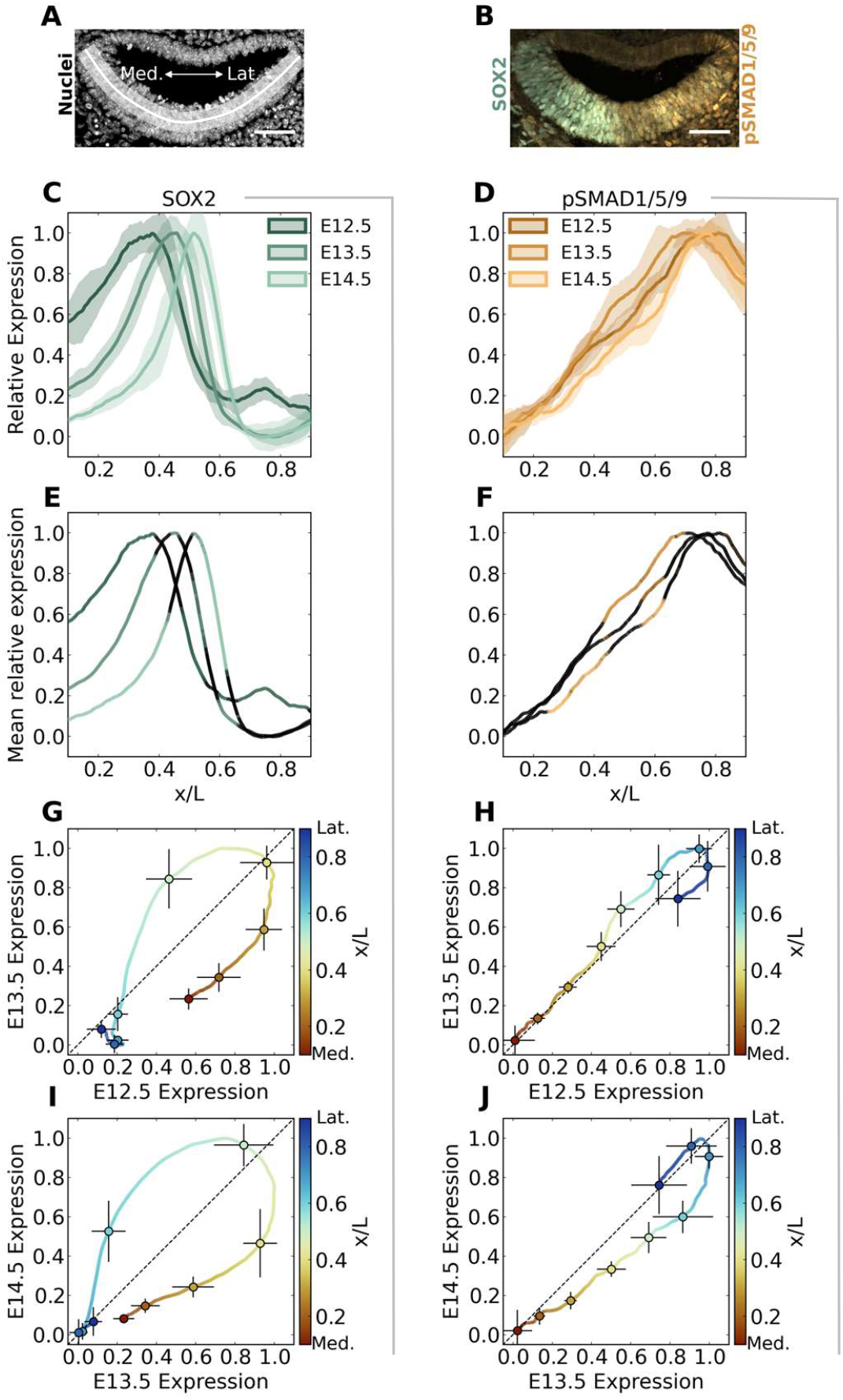
SOX2 is dynamic, while pSMAD1/5/9 is stable before the onset of differentiation. **(A-B)** Exemplar images from E12.5. Scale bar: 50 μm. **(A)** Profile data in the prosensory domain (PSD) is extracted for each channel from a region indicated by the curved white line, where x=0 at the medial (M) most point extending to the lateral (L) most point. **(B)** Medially concentrated SOX2(green) and laterally concentrated pSMAD1/5/9 (orange) are illustrated. **(C-D)** The final quantification results of SOX2 (C) and pSMAD1/5/9 (D) with mean profiles (bold) and standard deviation (shading) taken across N=15, 16, and 8 cochleae for E12.5, E13.5, and E14.5, respectively. **(E-F)** The colored portions of the profiles indicate signals significantly different from profile distributions from other stages (*p* → 0) calculated by an approximate two-sample Kolmogorov-Smirnov test. Black indicates where *p* ≥ 0.05. **(G-J)** Comparing SOX2 (G, I) and pSMAD1/5/9 (H, J) profiles between subsequent timepoints further illustrates the dynamic SOX2 signal (signals stray widely from the diagonal) compared to the stability of pSMAD1/5/9 (signals remain near the diagonal). The color scale indicates the medial-lateral location of the relative signals. Error bars indicate standard deviation in the signal at the timepoint it was measured depending on the error bar’s direction for 9 points at *x*/*L* = 0.1 to 0.9 in increments of 0.1.

### SOX2 Signaling is Dynamic While BMP Signaling is Stable During Early Morphogenesis

Inspecting the profile data, we first notice a clear difference in the stability between SOX2 and pSMAD1/5/9 (**Figure 2C, D**). To investigate this, we compare the estimated probability distributions for expression at each position for each timepoint using a Kolmogorov-Smirnov two-sample test (**Figure 2E, F**). This produces a p-value predicting whether samples at one position for one timepoint might come from a distribution produced by samples of another timepoint at the same position and vice-versa. Significantly different profiles from either of the other timepoints (*p* ≥ 0.05) are colored, and insignificant differences fade to black for 0.05 >*p* → 0. We also plot the position-specific relative signal amplitudes between consecutive timepoints, illustrating the rate and direction of change between timepoints (**Figure 2G-J**).

In these plots, position is encoded by color, with warm colors indicating medial and cool colors lateral. A trajectory along the diagonal indicates equal expression at corresponding positions between timepoints. To be above the diagonal means that an expression value seen at one position earlier has increased. Below the diagonal indicates a decrease in expression level at a specific position. Horizontal and vertical error bars on samples spaced equally in physical space indicate standard deviations in expression levels at earlier and later timepoints, respectively. Comparing the signal trajectories for SOX2 in **Figure 2G** and **Figure 2I**, medial positions below the diagonal and lateral points above indicate the lateralization of the signal through time. The similarly shaped large loops indicate that the rate at which SOX2 shifts is consistent between timepoints. Whereas with pSMAD1/5/9, the trajectory gently wavers above and below the diagonal, steady by comparison.

The systematic lateral shift and refinement of SOX2 boundaries from E12.5 through E14.5 to occupy the central region on the floor of the duct (**Figure 2C, E)** will become the medial and lateral bounds of the OC sensory epithelium. This refinement of SOX2 occurs in relative and absolute terms due to dynamic morphogen cues and physical cell movement. As the cochlea undergoes convergent extension, the radial width of the PSD decreases. Within this shrinking domain, the width of the region containing SOX2-expressing cells narrows as measured by the full width at half-max of the mean SOX2 profile, changing from covering 38% (115 μm) of 303 ± 23 μm at E12.5 to 27% (69 μm) of 254 ± 15 μm at E13.5 to 19% (49 μm) of 256 ± 19 μm at E14.5. There is also a significant secondary SOX2 peak at x = 0.74 on E12.5, coinciding with the pSMAD1/5/9 peak at this location (**Figure 4A**). At E12.5, the entire floor of the duct shows some SOX2 expression, where its minimum lies outside the cropped PSD shown. At later timepoints, it drops to its minimum at around *x*/*L* ≈ 0.75. The lateral smaller peak is still present at these later time points and is visible in a small group of up to four nuclei at the lateral edge outside the cropped edge (**Figure S1E, I**). With pSMAD1/5/9, the profiles in the medial 40% and lateral 30% of the PSD at E12.5 and E13.5 are statistically indistinguishable (**Figure 2F**), and the lateral peak locations are maintained between 0.7 ≲ x/L ≲ 0.8 for each of the three timepoints. In the central region, the signal gently wavers (**Figure 2F, H, J**).

### Interpreting Positional Information Encoded by SOX2 and pSMAD1/5/9

We also notice that the profiles medial to the peak at E12.5 and E13.5 appear to be linear, which is an ideal shape for encoding positional information. Fitting this region to a line indicates high correlation to the data at these timepoints (**Figure 3A, B**). The result of a linear pSMAD1/5/9 profile is significant for the transduction of positional information because this shape retains a one-to-one relationship between signal expression and position. Under sufficiently low noise, it encodes the theoretical maximum amount of mutual information between expression and position^17^.

**Figure 3.**
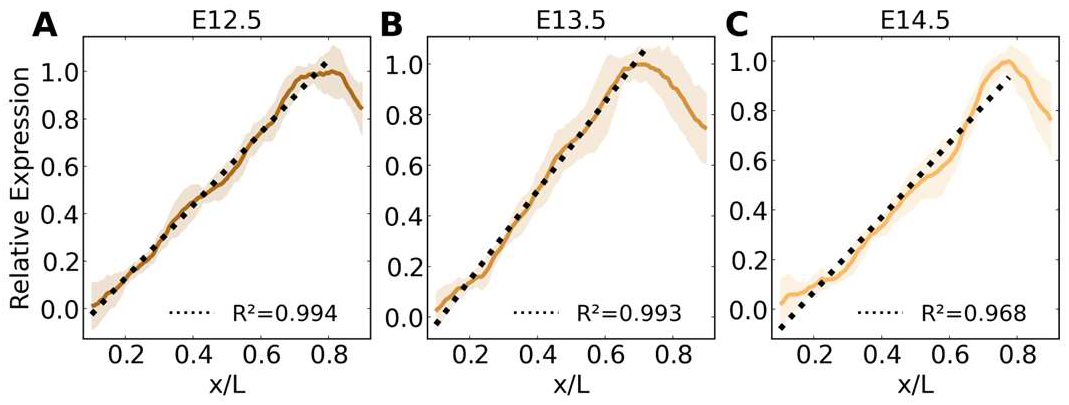
pSMAD1/5/9 profiles maintain a stable linear shape in the medial majority of the prosensory domain, especially during the E12.5-E13.5 period. pSMAD1/5/9 profiles (mean ± standard deviation) are strongly linear in the medial three-quarters of the prosensory domain at E12.5 (A) and E13.5 (B). By E14.5 (C), the linear shape begins to degrade. N = 15, 16, and 8 cochleae for E12.5, E13.5, and E14.5, respectively. R^2^: coefficient of determination for the linear fit (dotted line), which was fit over the region spanning the medial extreme and peak expression. Linear fit parameters for *y* = *mx* + *b* are *m*: 1.53, 1,76, 1.50 and *b*: -0.18, -0.21, -0.23.

To investigate the encoding of positional information for SOX2 and pSMAD1/5/9 individually and jointly, we used the decoding maps developed to predict the mapping between gap gene profile sets and space in *Drosophila*^18^. The SOX2 profiles individually reveal bimodal distributions in the medial half of the maps due to the steep rise and fall in this region (**Figure 4C, G**). However, when considered jointly with pSMAD1/5/9, SOX2 enables refinement of the decoding, although the lateral ‘smear’ still indicates that cells share a uniform identity in the region destined to become the outer sulcus (**Figure 4D, H**).

**Figure 4.**
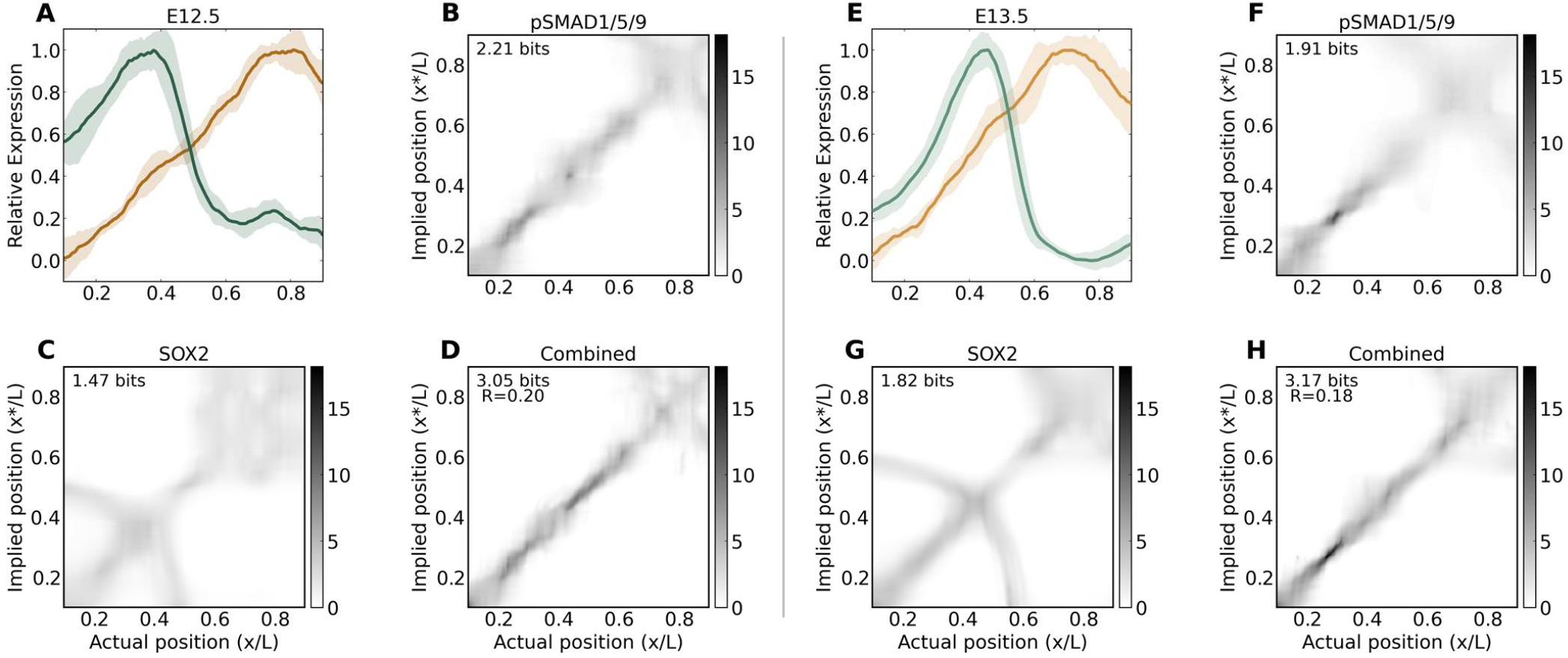
The mapping between pSMAD1/5/9 concentration and position in the medial two-thirds of the prosensory domain is 1:1 (along the diagonal of the maps), and the map is refined by SOX2. **(A-D)** E12.5. **(E-H)** E13.5. Map annotations indicate the estimated mutual information for individual and combined profile sets at each stage. The fractional redundancy R is indicated on combined maps to reveal the degree of independence for the information encoded in these two profiles. Map grayscales indicate probability density scaled against the single maximum value across all maps, individual and joint.

The mutual information encoded in these profiles was calculated to score the maps using the direct method also developed to analyze gap genes^17^, where the prior distribution *P*_*x*_(*x*) in the cochlea is considered as uniform by correcting for variations in nuclear density for each sample (see Methods). Inspecting the bit counts of the profiles, we see that there is approximately 20% redundancy in the mutual information encoded when considered jointly *I*({*g*_*i*_}; *x*) vs the sum of information estimated independently *I*(*g*_*i*_; *x*)^17^, calculated by the fractional redundancy:

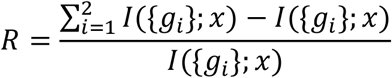

The joint mutual information at E12.5 of *I*({*pSMAD*1/5/9, *SOX2*}; *x*) ≈ 3.05 and at E13.5 of ≈ 3.17 bits indicates that these profiles contain sufficient information to designate 2^3.05^ ≈ 8 and 2^3.17^ ≈ 9 distinct features within the PSD at these timepoints.

However, interpreting bit counts directly into biological significance is difficult^20^. For a purely linear profile with uniform variance, it is calculated as 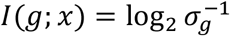, but with real data, the mutual information is dependent on a ratio between the spatially dependent slope and variance averaged over the domain. The variance of the pSMAD1/5/9 signal varies as a function of *x*/*L* along its region of linear fit (**Figure 3**).

The benefit gained by the linear pSMAD1/5/9 region in the medial 75% of the PSD can be demonstrated in that while the mean variance of SOX2 over this portion is higher than pSMAD1/5/9 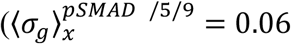 and 0.07, and 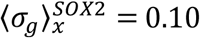 and 0.09 at E12.5 and E13.5, respectively), the approximate individual profile mutual information is significantly higher for pSMAD1/5/9 than SOX2 (*I*(*pSMAD*; *x*) = 2.23, 2.12 and *I*(*SOX2*; *x*) = 1.44, 1.85 at E12.5 and E13.5, respectively). This pattern holds for the full PSD profile, though it is less pronounced due to the lateral peak and decline of pSMAD1/5/9.

### Modeling Mechanisms to Produce a Linear pSMAD1/5/9 Profile

We sought to investigate whether we could reproduce the measured pSMAD1/5/9 profile using a minimal set of components in a 1D finite difference model of BMP activity between E12.5 and E13.5 (see Methods). First and most simply, we consider the effect of *Bmp4* mRNA distribution. It has been shown that signaling activity downstream of a morphogen can be influenced by the prepattern of its mRNA source and is required in some cases to model morphogen activity^31^. In the cochlea, although *Bmp4* transcription is constrained to a small lateral region, it is not concentrated into a single point source on E12.5 and exhibits nonzero expression between 0.4 < x/L < 1.0 (**Figure S4**). Considering only the transcript prepattern, simulations match pSMAD1/5/9 data poorly, however. The best fit solution is exponentially decaying in shape and has a lateralized peak compared to the data (**Figure 5A**).

**Figure 5.**
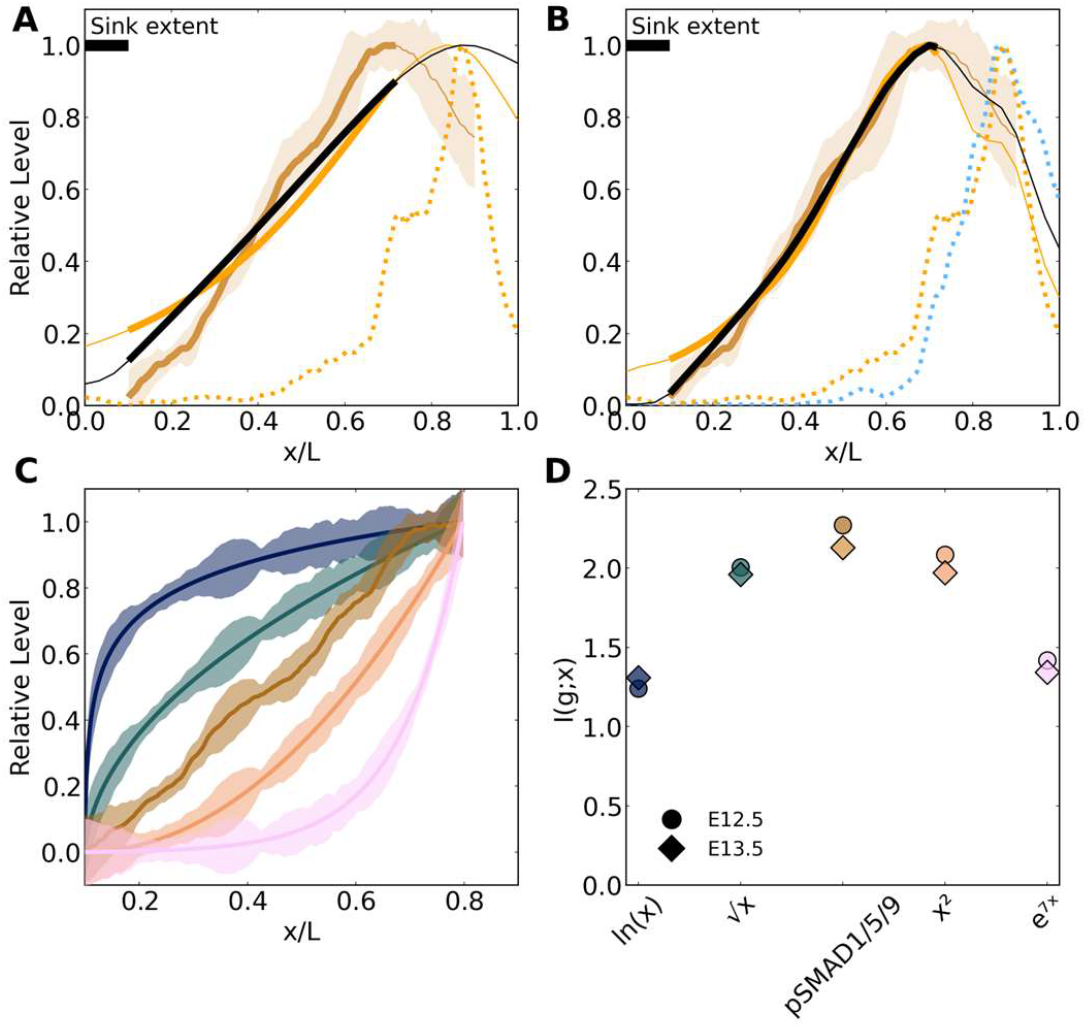
1D diffusion simulations suggest that the presence of a medial inhibitor of BMP signaling may help form a linear gradient, which optimizes positional information across the radial axis. The best-performing simulations are presented by minimizing the RMSE over the sampled parameter space on the linear pSMAD1/5/9 region (medial extreme to pSMAD1/5/9 peak). (A,B) Gold with shaded error represents E13.5 pSDMAD1/5/9, orange represents optimal transcript controlled (no-sink) solutions, black represents optimal solutions when a sink is considered, dotted orange represents the *Bmp4* mRNA distribution, and dotted blue represents the *Fst* mRNA distribution. Thick curves represent where the RMSE was calculated. (**A)** Transcription-controlled patterning with synthesis from a transcriptional source of *Bmp4* mRNA (orange dotted) and diffusion and degradation of BMP ligands only produces an exponential solution (orange smooth). When a hypothetical sink extending to *x*/*L* = 0.1 is added, the solution straightens (black), but a systematic error is still present due to the misalignment between the pSMAD1/5/9 (gold) and *Bmp4* expression peaks. **(B)** Assuming that FST binds BMP (uncertain in this context) and is produced proportionally to its mRNA distribution (blue dotted), a closer match is achievable. With no sink, the exponential shape is still present (gold). With the sink, data and simulation results match closely (black). **(C)** Measured E12.5 pSMAD1/5/9 data is shown in comparison to alternative shapes connecting the medial minimum to lateral maximum of expression. For synthetic profiles, the residuals between individual pSMAD1/5/9 measurements and the sample mean are mapped to the new shape for E12.5 (shown) and E13.5 (not shown). **(D)** Mutual information is maximized by the linear pSMAD1/5/9 at E12.5 and E13.5 individually compared to alternative shapes when accounting for measured variance.

To achieve a linear profile of pSMAD1/5/9 expression from a diffusive BMP4 source, additional mechanisms must be present. A linear gradient extending from a spatially constricted morphogen source was first proposed in 1970 by Francis Crick^3^. This proposition hypothesized the existence of a point-source of morphogen flux at one end of a domain, free diffusion of the morphogen across the domain, and a perfect point-sink at the opposite end where the concentration is fixed to zero. The steady-state solution to this system—regardless of initial conditions or the diffusion parameter—is a linear profile between zero at the sink and a constant concentration at the source, which depends on the source production rate. As theoretically simple as generating a linear gradient is, it is not a solution that has been observed in nature over any substantial portion of a patterning domain. Including a sink in the simulation—modeled as a hypothetical molecule to which BMP binds irreversibly and is present in excess and extends to *x*/*L* = 0.1—a linear profile is achievable (**Figure 5A**). However, there is still a significant error between simulations and data due to the offset between the *Bmp4* mRNA peak and the pSMAD1/5/9 data.

To account for this, we explore a plausible mechanism that could shift the BMP activity profile medially from the peak of *Bmp4* expression. It has been reported that Follistatin (FST), a secreted inhibitor of some transforming growth factor (TGF)-β ligands, is present in the lateral portion of the PSD at E13.5 and E14.5^32,33^. The *Fst* gene forms three isoforms with varying affinities to different TGF-β ligands, but all strongly bind Activin A with a strong preference for BMPs^34^. In postnatal cochlear cells extracted and dissociated to form organoids, doxycycline-induced FST-288 overexpression is followed by decreased expression of pSMAD1/5/9 and BMP reporters^35^. The same transgenic mice overexpressing FST during a more comparable stage of embryonic development did not show a difference in pSMAD1/5/9 *in vivo* when induced at E11.5 and measured at E14.5^33^. However, the spatial context of pSMAD1/5/9 signaling was not analyzed. Therefore, whether endogenously expressed FST interacts with BMP in this developmental context of the cochlea remains unclear. Assuming an interaction enables the model to perform well by shifting the simulated peak to the correct location (**Figure 5B**).

## Discussion

The mature OC is an extraordinary example of reproducibility and precision in cellular patterning that is established by dynamic phases of gene regulation coupled to and enhanced by mechanical remodeling. Our contribution reveals new results in the early morphogenetic activity of this process by quantitatively characterizing snapshots of crucial transcription factor profiles providing positional information along the radial axis of the PSD.

Analyzing the profile shapes of SOX2 and pSMAD1/5/9 at E12.5, E13.5, and E14.5, we observe significant differences in the stability exhibited over time between these molecules.

We note the production of a linear gradient for pSMAD1/5/9, which is an ideal shape for encoding positional information. The significance of this linear profile can be seen in its spatial decoding capabilities. The pSMAD1/5/9 profile provides a diagonal decoding map between implied and actual positions, where local precision is refined when considered jointly with the steeply ascending and descending SOX2 profile. Although there is positional error along this joint map’s diagonal (indicated by the vertical width of the probability density at a given actual position), all cells in this linear pSMAD1/5/9 region can detect their position only with inaccuracies regarding their nearest neighbors. That is, there are no bimodal ambiguities as with SOX2-only maps. The presence of minor imperfections in the positional specification of early differentiating HCs is supported by recent results demonstrating a patterning sequence where neighboring cells show some uncertainty in their final positions and are precisely refined into place by mechanical forces^14^.

The stability of the BMP-controlled signal compared to the dynamic shifting seen with SOX2 may indicate that pSMAD1/5/9 serves as a global reference for radial position across the early days of development, while SOX2 responds dynamically to transitioning Notch signals^4,36^. This stability can be seen in the diagonal decoding maps medial to the future OS across time (**Figure 2H, J**; **Figure S3B, E, H, K, N**) compared to the steadily shifting SOX2 signal (**Figure 2G, I**; **Figure S3A, D, G, J, M**). When interpreting the decoding maps, it has been shown in *Drosophila* that using a map derived from a single 5-minute window where mutual information is maximized, predictions for features of target genes can be reliably made at earlier, matching, and later timepoints^18^. In that system, the dynamic range of individual genes varies between ∼80% and 120% of the range used to construct the map. In the cochlea, this maximal information during phase one appears to occur at E13.5. However, the timescale considered here is on the order of days rather than minutes. The minimum and maximum absolute amplitudes of SOX2 and pSMAD1/5/9 (BMP) concentrations over this period may change significantly, making decoding from a single snapshot a disadvantage with the signal’s dynamic range changing substantially between days. The effect of BMP4 on pSMAD1/5/9 and IHC formation is sensitive to somewhere between 5 ng/mL^1^ or 10 ng/mL^35^ to 50 ng/mL, though exact endogenous concentrations are not known. Improved experimental techniques such as simultaneous imaging of endogenous reporter elements may help determine the effect of absolute concentration on decoding abilities.

Inspecting the decoding maps and their changes between consecutive days more closely, we note the emergence of a sharp increase in precision of the joint map around 0.25 ≲ *x*/*L* ≲ 0.30 between E12.5 and E13.5 (**Figure 4D, H**; **Figure S3C, F, I**). This range corresponds to roughly three nuclear widths, which could represent a radial reference point that anchors signals to this position for the next phase. By E14.5, when the HC-SC mosaic organization begins, there is an increased ambiguity around *x*/*L* = 0.5 with a sharp region medial to this point, which we speculate may represent lateral and medial regions of the emerging OC, respectively.

Mechanistically, the linear pSMAD1/5/9 profile is a surprising result for a readout from a diffusive morphogen which would typically fit an exponentially decaying curve. We model a series of plausible mechanisms that could produce such a shape, paying particular attention to recreating the linear aspect. The first and simplest mechanism accounts only for diffusion and degradation from the distributed production region set by the *Bmp4* transcript profile in the lateral portion of the PSD. However, it cannot replicate the linearity of the pSMAD1/5/9 profile. We next postulate a medial sink that removes BMP over a region of the medial PSD. In this instance, a linear profile is achieved, but the peak of BMP activity is misaligned with the observed pSMAD1/5/9 peak, indicating that more details in the BMP regulatory network must be investigated. FST is expressed in a convenient location to serve as a means to shift the BMP activity peak medially. Comparing quantitative radial pSMAD1/5/9 profiles between sections of FST-induced mice and or conditional knockouts with wild-type controls could sensitively detect relevant differences in BMP activity caused by excess FST.

The combined speculative mechanisms of lateral FST binding with BMP and an unidentified medial sink together closely match measured pSMAD1/5/9 data. Adding the sink enables the profile to achieve a linear shape. There are dozens of molecules known to interact with BMPs extracellularly^37^, though none are apparent candidates in the cochlea acting over E12.5-E13.5 from data available in the literature. The results presented here indicate that a systematic screen should be conducted over these candidates to sensitively detect their presence and localization in the PSD at these timepoints with fluorescent probes, which would better inform a mechanistic model. If a medial BMP sink molecule is identified, further studies may induce perturbations to disrupt the linear pSMAD1/5/9 gradient and better parameterize the model. An advantage of a medial sink is its relative insensitivity to parameter fine-tuning. If it is constrained to a limited lateral extent and sufficiently strong, it can effectively contribute to the linear gradient formation.

Additional unexplored mechanisms with increasing complexity are also feasible. For instance, BMP heterodimers promote higher pSMAD1/5/9 activity than homodimers of either ligand^38,39^. Both BMP4 and BMP7 are present in the PSD during the timepoints studied. While the simulations explored here are agnostic to the form of dimerization, local concentrations of either BMP7 or BMP4 monomers may be rate-limiting in the proportion of each dimer subset locally produced. The relative reaction rates of these diverse dimers coupled with the spatial profile of each ligand’s production domain may account for the linear profile in a more advanced version of the no-sink model presented here. Furthermore, sensitive radial measurements of other BMP pathway components such as type I and II receptors and intracellular cofactors such as SMAD4 should be considered to replace assumptions of their uniformity and sufficiency with data.

The subject of mechanisms forming morphogen gradients in diverse systems has been reviewed extensively (a sample of such reviews^40–46^). In *Drosophila* development, the transport of the BMP homolog Dpp is shown to fit a power-law curve from a source along the dorsal midline^47^ and exponential decay in the wing imaginal disc^48^. BMP is also exponential in the mouse neural tube^21^, zebrafish pectoral fin^49^, and the dorsal-ventral axis of the zebrafish blastula, which has even been shown to be under the influence of a source-sink mechanism with Chordin^50,51^. In fact, anywhere BMP and its homologs are documented to influence patterning, they manifest their signaling through an exponential (including sigmoidal) or power-law gradient. This is also true of other profiles under the control of secreted ligands such as Fgf8^52^, Shh^21^, Bicoid^53^, dpERK^54^, and Dorsal^55^. Though a linear fit may be approximated over a small region for any curve, a stable linear gradient exceeding a span of 200 μm, as pSMAD1/5/9 shows in the cochlea has yet to be reported.

But what if this profile had a different shape? Since mutual information is a function of profile shape and position-dependent noise in equal measure, we mapped the residual values between the pSMAD1/5/9 sample profiles and mean profile across the radial domain at E12.5 and E13.5 onto alternative profile shapes (E12.5 shown in **Figure 5C**) and calculated the corresponding mutual information (**Figure 5D**). Mathematically, a linear profile is predicted to encode maximum information compared to alternative shapes when noise is sufficiently low^17^. Here, the linear pSMAD1/5/9 profile indeed outperforms nonlinear profiles. Both experimental downregulation and upregulation of BMP severely perturbs radial patterning^1,23^, indicating that every bit of this information is needed to properly position target gene expression and thus subsequent domain boundary formation.

Furthermore, although this is the best-resolved quantitative signaling data in cochlea development to date, another recent advance in a study of positional information in neural tube development shows that profile precision may be significantly sharper than what is possible to detect with the standard conjugated antibody fluorescence methods used here compared to readouts from conjugated direct morphogen reporters^22^, even when experimentally-imposed variability is corrected for by model-based normalization^17,56^.

This underscores the importance of calibrating our interpretation of positional information between the optimal decoding assumption of an upper limit to positional mapping. Decoding is sensitive to the (im)precision of measurements, but when measurements are close to exact, complex gene regulatory mechanisms downstream of nuclear transcription factor concentrations may act as bottlenecks for this information transmission^19,57^. Even so, the quantitative results here provide a basis of helpful interpretation for the sensitivity to pattern disruption when BMP or SOX2 signaling are experimentally perturbed during these stages^1,23,25,26^, and such decoding maps may be used to predict features of target gene expression profiles such as *Atoh1* and *Id1, Id2*, and *Id3*^4,18^.

Primarily, our results demonstrate the unique presence of a linear gradient resulting from a diffusive morphogen system, establishing the groundwork of patterning precision in the cochlear PSD. This reveals a rich opportunity for further study of what mechanisms are at play to control such a process, and the informatics tools employed provide a new lens for interpreting patterning in this system.

## Methods

### Experimental

Swiss Webster mice (Charles River Crl:CFW(SW)) were mated, and pregnancies were timed starting at E0.5 on the day a plug was observed to collect embryos at E12.5, E13.5, and E14.5. Animal procedures were conducted following protocols set by the Purdue Animal Care and Use Committee (PACUC).

Profiles at each stage were acquired from mid-basal cochlear cross sections for uniformity in developmental maturity since differentiation occurs in a base-to-apex direction along the longitudinal axis, while cell cycle exit in the sensory domain occurs in the opposite direction^58^.

### Immunofluorescence Staining

20 μm sections were stained similarly to a previous report^23^. The sections were first incubated in Trypsin (0.5%) for 45s and immediately washed gently three times with PBS with 0.1% Triton X-100 (PBS-T). This was to eliminate nonspecific granulation of the primary antibody in the tissue. Trypsin digestion in this manner at this high concentration and short exposure helped to clear components that caused the artifacts. Following washes, sections were fixed in PFA (4%) for 15min, washed and permeabilized with PBS-T three times for 5min each, and blocked with donkey serum (GeneTex #GTX27475 Lot 822003572; diluted to 2%) for 1h. Sections were incubated overnight at 4°C in a primary antibody solution consisting of rabbit-anti-pSMAD1/5/9 (1:600, Cell Signaling Technology #13820 Lot 3) and goat-anti-SOX2 (1:400, R&D Systems #AF2018 Lot KOYO418121) in PBS-T with 2% DS. The next day, sections were moved to room temperature and washed three times for 5min each with PBS-T. Secondary antibodies (Thermo Fisher Scientific donkey anti-rabbit Alexa Fluor 488 #A-21206 Lot 2156521 and donkey anti-goat Alexa Fluor 568 #A-11057 Lot 2160061) were mixed to 1:500 in PBS-T with 2% DS and added to sections to incubate for 1h. Nuclear staining with TO-PRO-3 (Thermo Fisher Scientific #T3605, 1:2500 in PBS-T with 2% DS).

### RNAscope Staining

The mRNA for *Bmp4* and *Fst* was stained using the ACD RNAscope Multiplex Fluorescent Reagent Kit v2 protocol for fixed frozen tissue. The probes used were Mm-Bmp4-O1-C3 (#527501-C3) and Mm-Fst-O3 (553211), which were developed using dyes by Akoya Biosciences, including Opal 520 (#FP1487001KT) and Opal 690 (#FP1497001KT).

### Imaging

After mounting and drying coverslips, confocal imaging was conducted with a Zeiss LSM 800 using a 20X water-immersion objective for immunofluorescence and a 40X oil-immersion objective for RNAscope. Briefly, laser power and gain were adjusted for each image to achieve maximum brightness with no pixel saturation in the cochlea. Samples were scanned one channel at a time with pinhole diameter set to the optimal value for each channel at a resolution of 1024×1024 with16-bit pixel depth and averaging set to 2 to create a 1 μm increment z-stack through the full 20 μm section thickness.

## Computational

### Preprocessing

#### Manual preprocessing (FIJI^59^)

Images were oriented to place the floor of the duct on the bottom and the medial edge of the PSD on the left. A maximum intensity projection was applied to each channel across z-stacks. To extract linearized fluorescent data, a spline fit line was drawn across the middle of the floor of the duct from the medial edge to the lateral edge with a width of 80 pixels. Extracted data was saved in CSV format for further processing.

#### Automated preprocessing

The extract-transform-load (ETL) pipeline for generating quantified data used the following steps:

##### 1) Cancel noise added to the signal caused by variable nuclear density

The nuclear channel was used as a measurement for noise in the signal due to variable nuclear density. This can be canceled using noise regression, where a true signal *s*(*x*) is the difference between a signal measurement *m*(*x*) and noise *n*(*x*).

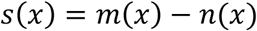

Using the nuclear channel as a reference measurement for the noise *n*_*ref*_(*x*) induced by variable nuclear density, we assume it has a linear relationship with the true noise term such that our best estimate for the noise 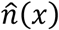 is:

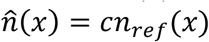

And our best estimate for the signal *ŝ*(*x*) is:

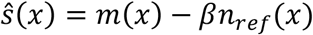

Identifying a *β* that minimizes the ‘energy’ in *ŝ* (*x*) given by ∑_*x*_ *ŝ* (*x*)^2^ will yield a suitable estimate such that:

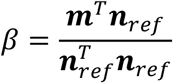

##### 2) Smooth profiles

A 20 μm moving average was applied.

##### 3) Resample

Data were resampled to a uniform size higher than the original using linear interpolation.

##### 4) Normalization and variance minimization

First, we anchor the minimum and maximum values of the mean for each signal to 0 and 1, respectively. Since the minimum value for pSMAD1/5/9 across all stages is near the medial edge of the PSD (and in the roof), this 0 corresponds with that minimum and is not necessarily zero concentration. Similarly, the lateral edge is where SOX2 is minimum at E12.5, though this appears to be close to zero concentration, as seen in the roof of the duct (see **Figure S1**). Assuming the variance between sample profiles was primarily due to experimental causes, we use model-based normalization^17,56^ to minimize this. For reference, *α* and *β* were identified for each embryo *i* to minimize the *χ*^2^ variance between each profile in the set and the mean, defined as:

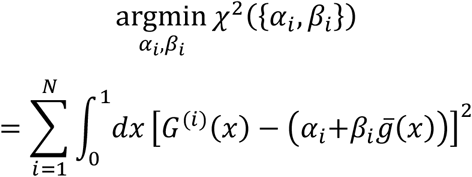

In the case of mesenchymal pSMAD1/5/9, the minimum of the mean profiles was not anchored to zero prior to variance minimization.

##### 5) Crop outer 10%

The data at the medial and lateral 10% for each profile were cropped to eliminate errors caused by ROI widths extending outside the cochlea in some samples.

#### Analyses

Positional information estimation for a gene set {*g*_*i*_} was implemented as mutual information using the “direct method” with settings for binning (Δ) and bootstrapping (*M*) as previously described^17^, reprinted here for reference.

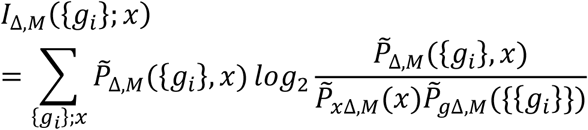

The decoding maps—presenting the probability density of an implied position *x*^*^ given an actual position 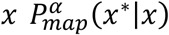 averaged over samples *α*— were implemented using the previously published method^18^, reprinted here for reference.

Single gene:

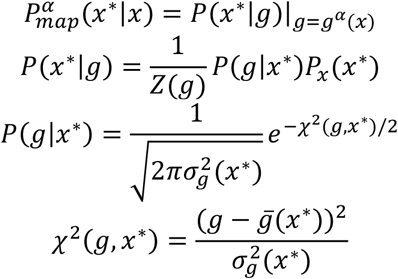

Two or more genes:

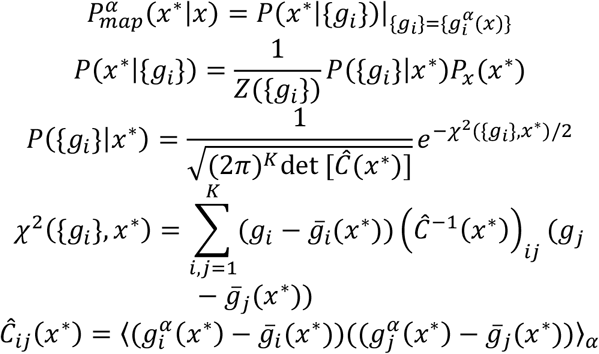

### Modeling

To inform the model with data, we set initial conditions to the mean pSMAD1/5/9 profile taken at E12.5. The model simulated 24 h of system evolution for a comparison to the mean pSMAD1/5/9 profile taken at E13.5. We performed a fluorescent RNAscope of *Bmp4* transcripts at E12.5 to create a spatial function representing the flux of BMP4 ligands into extracellular space and used the E12.5 pSMAD1/5/9 profile as the initial condition. Since the PSD is not assumed to be a closed volume (i.e., ligands may diffuse out from it into the surrounding mesenchyme), we extended the full simulation domain Ω beyond the ∼300 μm span of the PSD. We use Neumann (no-flux) boundary conditions (BCs) at *x* = 0 and *x* = L = 600 *μm* on this extended domain, where the PSD resides within the subdomain *ω* ∈ (150,450) *μm*, noting that the mesenchymal profiles of pSMAD1/5/9 decay exponentially and level off to a slope of 0 (**Figure S2**). A sink is modeled as a hypothetical molecule present in excess that irreversibly binds BMP over a series of binding affinities and extends from the medial edge of the PSD to a series of radial positions up to *x*/*L* = 0.5. The sink’s profile denoted as *S* (*x, b*), is modeled as either a unit-step (Heaviside) concentration function or a linear gradient decreasing from a maximum value at *x* = 0, where its domain extends from the medial extreme to a lateral boundary at *x*/*L* = *b* = (0.0,0.1), where *b* = 0 corresponds to the no-sink (i.e., transcript-control only) condition. The sink is assumed to exist in excess where it is expressed and irreversibly binds BMP with a parameterized affinity. Dissociation between BMP and the sink is neglected.

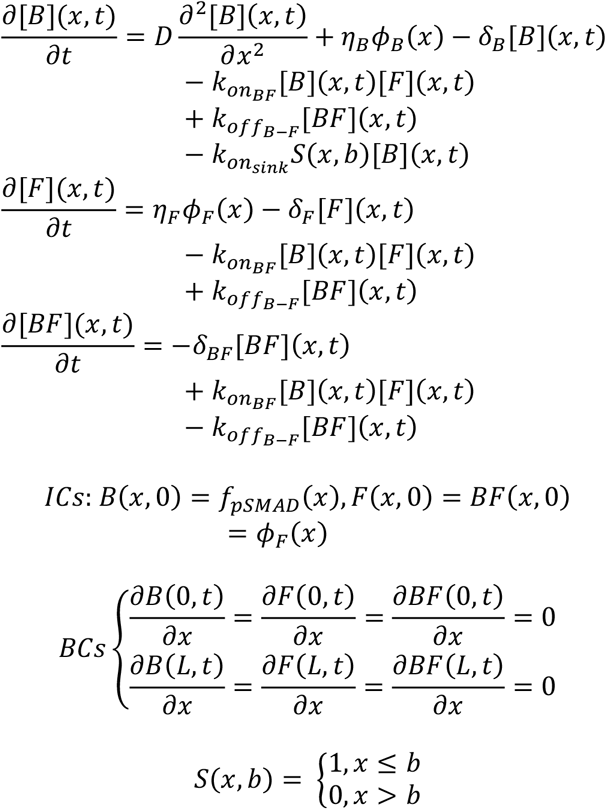

Here, [*B*](*x, t*) is the BMP ligand concentration [*nM*] which is assumed to be proportional to pSMAD1/5/9 level. Likewise, [*F*](*x, t*) and [*BF*](*x, t*) are the free FST and bound BMP-FST complexes. The synthesis profiles are given by *ϕ*_*B*_(*x*) and *ϕ*_*F*_(*x*)—the normalized E12.5 *Bmp4* and*Fst* profiles measured by RNAscope, respectively. The E12.5 initial condition of the mean pSMAD1/5/9 profile is *f*_*pSMAD*_(*x*).

For each condition, we performed a finite difference parameter optimization using a particle swarm with a population of 200 over 30 randomly seeded generations, iterating until the previous optimal solution was not improved. The cost function to minimize was the root-mean-squared error (RMSE) between [*B*](*x, T*) and the E13.5 pSMAD1/5/9 measurements. Solutions were penalized where the maximum amplitude of the simulation at the final timepoint exceeded that of the initial condition by more than 10x or decreased to less than 0.5x to avoid excessive fold-changes. Next, the solutions were normalized to a maximum of 1.0 since the true fold-change in expression concentration between E12.5 and E13.5 is unknown. This restricts our comparisons to overall shape rather than absolute quantities, which our semi-quantitative data does not permit across timepoints. The region within the PSD subdomain between the medial extreme and pSMAD1/5/9 peak was used in this calculation to emphasize the attempt to recreate a linear profile, depicted as the bold region of the curves in **Figure 5**.

Inspecting the parameter values for optimally chosen simulation solutions when FST is included (**Table 1**), we see that the diffusion rate used for BMP to transport from its lateral source to the pSMAD1/5/9 data declines when a sink is present. However, the diffusion rate is still on the order of 10-fold higher than measured biological diffusion^51^ around 4 μm^2^/s. When this parameter is fixed, we expected the BMP decay rate to drop substantially to allow it to transport medially (simulated profiles not shown). In each case, this is the pattern we observe by ∼10-fold or greater. Lastly, we note that the optimized dissociation constant for BMP4 and FST (*K*_D_ = *k*_*off*_/*k*_*on*_) when a sink is present comes out to 7.4 nM, a reasonable result comparing to 22.5 nM measured via surface plasmon resonance.

**Table 1.** BMP model parameters for optimal simulations. Parameter selections are included for models when FST binding with BMP is and is not considered, when a medial BMP sink is and is not considered, and when the BMP ligand diffusion constant is varied and is held to a commonly measured value of 4 μm^2^/s^*51*^. Optimized RMSE values indicate the best fit for the model using FST and a sink. Parameters are defined as follows: *D*:BMP ligand diffusion constant, *η*_*B*_: BMP ligand production rate, *η*_*F*_: FST production rate, 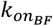: On rate constant for BMP binding with FST, 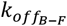: Off rate constant for BMP unbinding with FST, 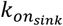: on rate for BMP and a hypothetical BMP sink molecule present in excess, *δ*_*B*_: decay rate of BMP ligand, *δ*_*F*_: decay rate of FST, *δ*_*BF*_: decay rate of bound BMP-FST. LB: lower bound, UB: upper bound.

| Param.<br>Symbol | With FST<br>Without sink |  | With FST<br>With sink |  | Without FST<br>Without sink |  | Without FST<br>With sink |  | LB | UB | Units |
| --- | --- | --- | --- | --- | --- | --- | --- | --- | --- | --- | --- |
|  | *Free*<br>*Diff.* | *Fixed*<br>*Diff.* | *Free*<br>*Diff.* | *Fixed*<br>*Diff.* | *Free*<br>*Diff.* | *Fixed*<br>*Diff.* | *Free*<br>*Diff.* | *Fixed*<br>*Diff.* |  |  |  |
| $D$ | 7.4E+1 | 4.0E+0 | 4.4E+1 | 4.0E+0 | 6.4E+1 | 4.0E+0 | 9.5E+1 | 4.0E+0 | $10^{-2}$ | $10^2$ | $\mu\text{m}^2/\text{s}$ |
| $\eta_B$ | 4.8E-1 | 3.6E-2 | 1.3E-1 | 1.0E-2 | 1.7E-1 | 1.0E-2 | 2.2E-1 | 1.0E-2 | $10^{-2}$ | $10^1$ | nM/s |
| $\eta_F$ | 2.8E0 | 3.2E+0 | 2.0E0 | 1.0E-2 | — | — | — | — | $10^{-2}$ | $10^1$ | nM/s |
| $k_{onBF}$ | 1.0E0 | 1.0E+0 | 4.3E-2 | 1.0E+0 | — | — | — | — | $10^{-4}$ | $10^0$ | 1/nMs |
| $k_{offB-F}$ | 1.0E0 | 3.1E-1 | 3.2E-1 | 4.3E-1 | — | — | — | — | $10^{-4}$ | $10^0$ | 1/s |
| $k_{on_{sink}}$ | — | — | 1.0E0 | 9.3E-3 | — | — | 1.6E-1 | 3.5E-3 | $10^{-4}$ | $10^0$ | 1/nMs |
| $\delta_B$ | 4.4E-2 | 5.3E-4 | 1.2E-2 | 1.0E-5 | 5.8E-3 | 3.6E-4 | 1.6E-3 | 2.1E-4 | $10^{-5}$ | $10^{-1}$ | 1/s |
| $\delta_F$ | 8.0E-2 | 5.2E-2 | 1.0E-1 | 4.1E-2 | — | — | — | — | $10^{-5}$ | $10^{-1}$ | 1/s |
| $\delta_{BF}$ | 1.9E-2 | 1.0E-5 | 5.5E-2 | 2.5E-3 | — | — | — | — | $10^{-5}$ | $10^{-1}$ | 1/s |
| RMSE | 0.048 | 0.085 | **0.026** | 0.094 | 0.119 | 0.119 | 0.091 | 0.096 |  |  |  |

Simulations were implemented using MethodOfLines.jl,whichrelies on ModelingToolkit.jl^60^ and DifferentialEquations.jl^61^.

## Acknowledgments

We thank Dr. Hari Bharadwaj of the Speech, Language, and Hearing Science department at Purdue University, Dr. Donna Fekete of the Department of Biological Sciences at Purdue University, Dr. Mariela Petkova of the Department of Molecular and Cellular Biology at Harvard University, and Dr. Joseph Zinski of the Department of Cell and Developmental Biology at the University of Pennsylvania for helpful discussions. This work has been supported by the National Institute on Deafness and Other Communication Disorders (NIDCD) T32DC016853, NIDCD R21DC016376 to V.M., and the National Institutes of Health (NIH) R01HD073156 and National Science Foundation (NSF) 2120200 to D.M.U.

## Author Contributions

M.J.T., D.M.U., and V.M. conceived the project and wrote the manuscript. M.J.T. performed the experiments and analyses.

### Data and Code Availability

All data generated for this publication and code used for processing, analyses, and figure generation are available in the following repository.

https://dagshub.com/thompsonmj/cochlea-positional-information

