## Supplemental Figures for "Early precision of radial patterning of the mouse cochlea is achieved by a linear BMP signaling gradient and is further refined by SOX2"

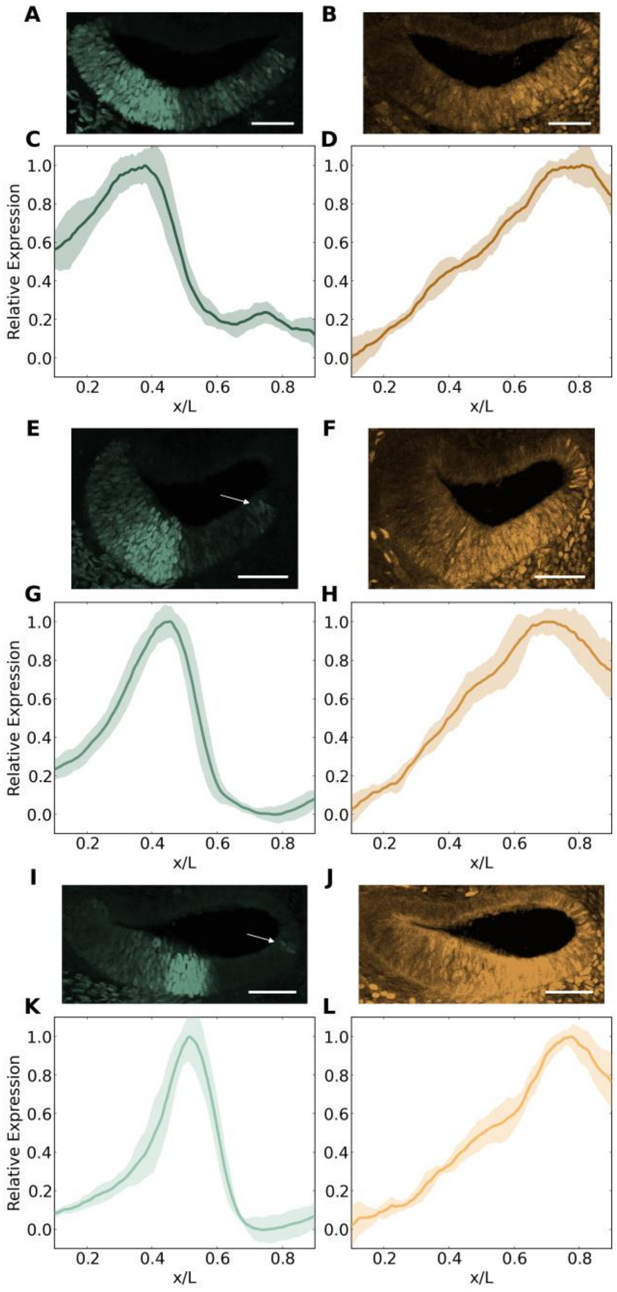


**Figure S1. Representative images of SOX2 and pSMAD1/5/9 at E12.5 (A-D, N=15), E13.5 (E-H, N=16), and E14.5(I-L, N=8).** SOX2 shows nonzero expression throughout the floor of the cochlear duct in the PSD at E12.5 (**A, C**)**.** The floor of the signal for pSMAD1/5/9 is taken at the medial extreme of the PSD, where it is minimal (**B, F, J**). Thus pSMAD1/5/9 is slightly above zero at $x = 0.1$ where it is cropped (**D, H, L**). There is a small group of $SOX^+$ cells near the lateral extreme of the duct floor at E13.5 and E14.5 (arrows, **E, I**), which is excluded from profiles due to cropping (**G, K**). Portions of images may appear saturated due to brightness scaling for clear presentation. Scale bars: 50 µm.

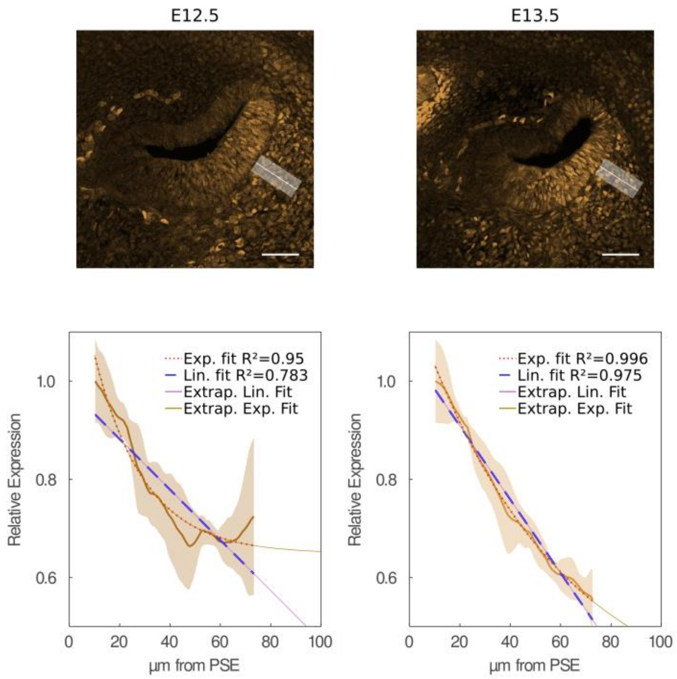


**Figure S2. The pSMAD1/5/9 profile in the mesenchyme near the peak best follows an exponentially decaying pattern.** Though the decay properties vary between E12.5 and E13.5, the exponential shape is consistent with simple synthesis, diffusion, and degradation of BMP ligand in this surrounding tissue compared to the linear profile seen in the prosensory domain. N=9, 8 for E12.5 and E13.5, respectively. Scale bars: 50 µm.

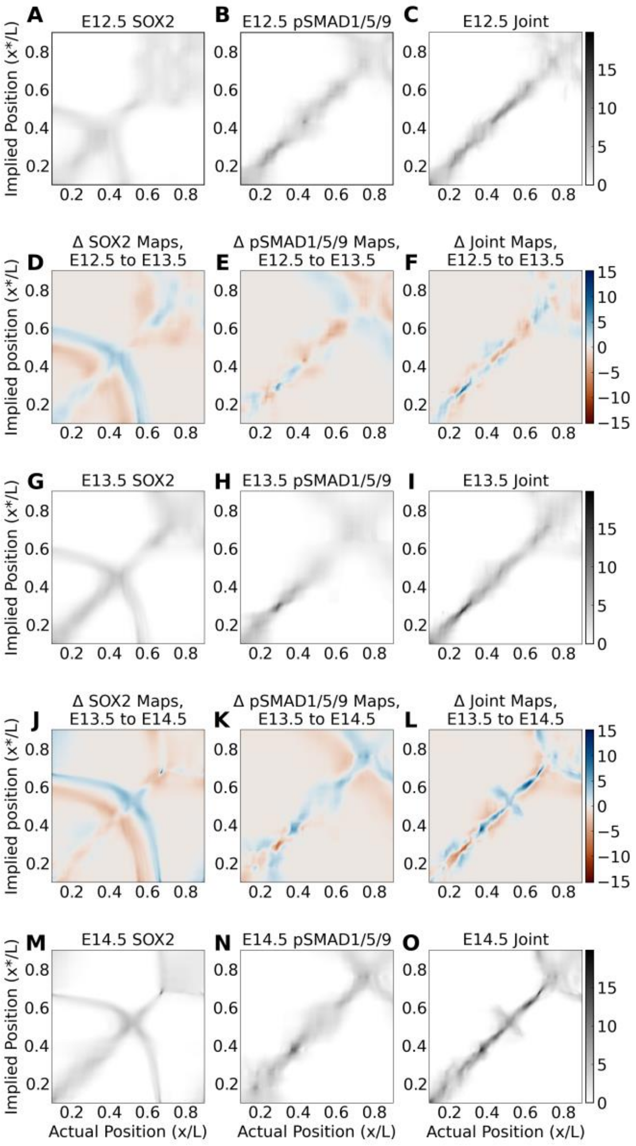


**Figure S3**. **Joint decoding maps maintain peaks close to the diagonal along the medial ~2/3 of the prosensory domain across E12.5, E13.5, and E14.5.** Grayscale maps are decoding probability density snapshots at each time point for individual (first and second columns) and joint (third column) profiles. Each of these is scaled against the single maximum across all maps at all timepoints to aid visual inspection. Each colored map shows the change in probability density between the map above and below itself, highlighting the dynamics between days. These maps are also scaled against the single maximum across all difference maps. In the joint difference map from E12.5 to E13.5, the map sharpens in the medial portion, broadens in the center, and changes minimally in the lateral portion of the PSD. A notable increase in precision between these snapshots occurs between x=0.25 and x=0.30, corresponding to roughly three nuclear widths. Between E13.5 and E14.5, the lateral ambiguity remains, but the sharpest positions flank the central ~10% of the domain.

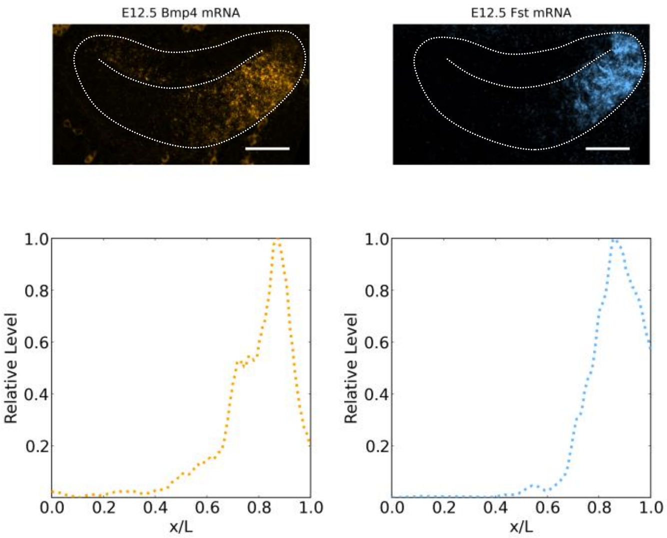


**Figure S4. E12.5 Bmp4 and Fst mRNA co-stained fluorescent RNAscope.** Used as the profile for the corresponding source flux terms in BMP simulations. Related to . Scale bar: 50 µm.
